# Stable isotope analysis (*δ*^13^C, *δ*^15^N, *δ*^34^S) reveal divers manuring practices of rye (*Secale cereale*) in northern Europe since 1500 years

**DOI:** 10.1101/2024.08.12.607590

**Authors:** Frank Schlütz, Felix Bittmann, Susanne Jahns, Sonja König, Lyudmila Shumilovskikh, Michael Baumecker, Wiebke Kirleis

## Abstract

First analysis of stable isotopes in rye from northern and eastern Germany provides insights into the early history of rye cultivation from the migration to the late medieval period. A comparison of the *δ*^15^N with modern experiments reveals that already the early cultivation involved the practice of intensive manuring. The intensity of manuring does not demonstrate a discernible trend over time but high variability. In some instances, it is likely that rye was cultivated as part of a rotation cultivation system including manuring. Through its *δ*^34^S values some rye turned out to be manured with marine peats. Rye, reputed to be a frugal crop, was cultivated on poor sandy soils to a limited extent only. Statistically significant correlation between the ^13^C content and yields demonstrates that the highest yield success on dwelling mounds was achieved, while the majority of fields on dry sandy soils yielded in relation only 40 to 70 %. Manured rye facilitated the emergence of multi-headed village communities and towns and stabilized the political power systems. The cultivation on mounds flooded by winter storm surges emphasises that summer rye was initially cultivated, before winter-rye became the dominating crop in Germany until the mid-20th century.

## Introduction

Rye was only occasionally and temporarily cultivated in the Levante during the Aceramic Neolithic or even earlier (Hillmann 1978, Hillmann et al. 2001) [1,2]. It came to Europe as an arable weed infesting the grain fields of barley and wheat species in the early Neolithic (Behre 1992, Schreiber et al. 2021) [3,4]. The number of archaeobotanical finds increase there with the Bronze Age, but it was not until the Iron Age that first rye-dominated finds indicate the intentional cultivation as a staple food (Schultze-Motel 1988, Wasylikowa 1991, Behre 1992) [3,5,6].

This post-Neolithic domestication took place independently at various places in Europe and at the northernmost rim of SW-Asia within the centuries around the turn of the Christian era (Seabra et al. 2023, Schultze-Motel 1988, Gyulai 2014, Behre 1992) [3,5,7,8]. The domestication of rye from a weed admixture to a cereal in its own right, was discussed by Behre (1992) [3] against the background of the low soil and climate dependency of rye in conjunction with climate deterioration and the transition to harvesting close to the ground, which became possible with iron sickles (Behre 1992) [3].

In the sandy areas of northern Germany, a work-intensive permanent cultivation of rye (‘Ewiger Roggenbau’, eternal rye) developed with the start of the second millennium CE. This so called ‘Plaggenwirtschaft’ comprised the cut of heath sods as stable bedding and depositing the resulting soil-dung mixture as plaggen manure on certain plots for a permanent field cultivation. The surface of the resulting plaggen-soils arose above the surrounding landscape and the steady influx of nutrients allowed a permanent rye cultivation including the harvesting of the straw. Only about every 10 or more years a one-year fallow phase was necessary (Conry 1974; Behre 2000; Beck et al. 2010) [9–11].

Compared to the natural baseline, nitrogen concentration and the heavy nitrogen isotope ^15^N are raised in dung. Consequently, cereals cultivated on manured soils can exhibit remarkable high *δ*^15^N values. Despite the sparse research on isotopes from rye of archaeological contexts, charred rye grains with relatively high ^15^N content are known for southern Scandinavia already from the 5^th^ to 6^th^ century CE (Larsson et al. 2019) [12], and are therefore half a millennium older than the medieval intensive plaggen farming system of rye (Beck et al. 1989) [11]. We here try to elucidate the early north European cultivation of rye including new isotope measurements of ^13^C, ^15^N and ^34^S on rye from settlements of eastern Germany and the North Sea coast, dating into the migration period, Slavic period and medieval times, as well as the ^15^N from manured and unmanured cultivation experiments. Together with isotope data published so far (Larsson et al. 2019; Brinkkemper et al. 2017, Hamerow et al. 2020, Golea et al. 2023) [12–15], this gives a fairly clear idea of how manuring made rye one of the dominant crops in parts of northern Europe for over 1500 years until the mid-20th century (Büttner et al. 2001) [16].

## Material

From two agricultural experiment sites in eastern Germany *δ*^15^N values have been measured from rye grains of four different manuring levels harvested in 2017, from each plot 20 grains. Harvesting included also the straw, resulting in a high nutrient extraction. At Thyrow, 30 km south-west of Berlin, the *Static Nutrient Deficiency Experiment* on slightly silty sands with low organic content started in 1937. The Albic Luvisol developed from rust-coloured forest soils, is of low water-holding capacity and a typical soil of the moraine landscape of eastern Germany. The annual mean temperature and precipitation are about 9.7°C and 533 mm (1991-2020) respectively, with water deficit in the summer month. The cultivation of winter rye as monoculture without rotation started in 1998. It includes plots unmanured and manured with dung (15t/ha*a) (Baumecker 2018) [17]. From both, *δ*^15^N of 20 individual rye grains from the yield 2017was measured. The *Eternal Rye Experiment* at Halle/Saale started in 1878. Winter rye is cultivated since then in monoculture without crop rotation. The base of morainic material is here covered by sandy loess, from which a Luvic Chernosem developed (Böttcher et al. 2000) [18]. Mean annual temperature and precipitation are 9.6°C and 516 mm (1981-2010) respectively. An unmanured plot was used for marker experiments with enriched ^15^N (Schliephake et al. 1999) [19] and is therefore neglected here. From the yield 2017 of two manured fields, 20 grains of each were measured for *δ*^15^N. One field is manured since 1878 by 12t/ha*a stable dung, the other by the same amount from 1893-1952 but not thereafter anymore (Böttcher et al. 2000, Herbst 2017) [18,20].

The analysed charred cereal grains derive from archaeological excavations conducted during the last decades in northern Germany, in the Lower Saxony coastal area of the North Sea and in the state Brandenburg, eastern Germany. In addition, isotopes on rye from Georgia were measured. The rye from the coastal area originates from two geological and hydrological different landscapes - the wet marshlands of the German clay district and the dry Geest (upland). While the subsoil of the flat clay district consists mostly of peats and clays formed during the Holocene, the hilly Geest is characterised by sands and gravels from the Saalian (or penultimate) glaciation.

In the German clay district (marshland, ‘Marsch’) people lived during phases of low marine influence on flat, dry fallen ground or artificial dwelling mounds - terps in Dutch or ‘Wurten/Warften’ in German, erected on the dry fallen ground. The houses were divided into a living space and a stable part (Brandt 2002) [22]. Manure, drifted organic material and marine clay were stacked in alternating layers to raise the dwelling mounds following sea level rise and small mounds with single or few houses often fused to lager village mounds (Schmidt 1994) [23]. The top of the Wurt Niens is today in 3-4 m asl and occupied by a farm, that of Oldorf (up to 5 m asl) carries a village. Niens was founded on an embankment wall of a former tideway surrounded by dry fallen marine sediments in the north of the today’s peninsula Butjadingen (Fig. 1) (Brandt 2002) [22]. The early medieval colonization started on level ground at the end of the 7^th^ century AD and ended in the first half of the 9^th^ century. At the beginning agriculture and livestock keeping took place under freshwater conditions, but with the rising sea level soon a dwelling mound was erected as protection against winterly storm tides (Brandt 2002, Behre 1991) [22,24]. Plow tracks and traces of cattle from the initial colonization are preserved below parts of the mound giving evidence of the economic activities. The remains of rye analysed dates into the 7^th^-8^th^ century. As one of the oldest Frisian settlements, Oldorf was founded in the first half of the 7^th^ century. As in Niens, the economy was mainly based on livestock breeding, and the mound also partially covers former arable land. The medieval settlement ended in the 11^th^ century (Brandt 2002) [22]. Rye found here is ^14^C-dated into the 9^th^ century.

**Fig. 1.**
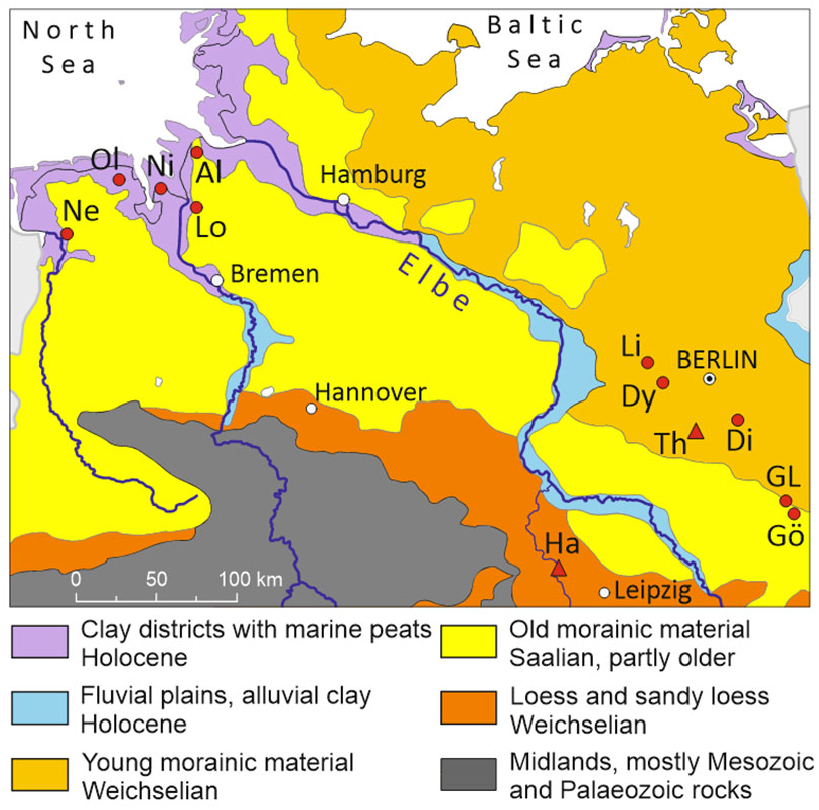
Soil regions of northern Germany with the sites Neermoor (Ne), Oldorf (Ol), Niens (Ni), Loxstedt (Lo), Altenwalde (Al), Lietzow (Li), Dyrotz (Dy), Diepensee (Di), Groß Lübbenau (GL), Göritz (Gö), Thyrow (Th), Halle (Ha) (map source BGR 2023 [21]).

Loxstedt and Altenwalde are settlements situated on the Geest ridge Hohe Lieth (about 25 m asl at Altenwalde, surrounded by the clay district to the north, west and east). Loxstedt, south of Bremerhaven, was an agricultural village of more than a dozen long houses with stables and about 100 pit houses of the 4^th^-6^th^ century (Zimmermann 2001) [25]. The archaeobotanical material of Altenwalde including rye was found during an emergency excavation in several pits. In some of them pottery of the 14^th^ century occurred (Behre 1998) [26]. Barley from the same context was dated to the 13^th^-14^th^ century. The remains of rye from Neermoor are from a late-medieval castle built at the lower part of a flat Geest slope in the 13^th^ century and destroyed not later than AD 1409. The excavated castle measured 70 by 70 m and was surrounded by a 6 to 8 m wide water-filled moat (Hüser 2015, Kegler and König 2015) [27,28]. Adjoining to the west large areas of raised bog *Sphagnum*-peats covered by brackish clays are situated (NIBIS) [29].

The material from eastern Germany derives from five excavations spread over the county of Brandenburg, including sites around Berlin (Jahns et al. 2018) [30]. The geological subsoil is dominated by morainic sediments of the last and second last glaciation (LBGR). While the older material from the penultimate Saalian glaciation was leached over time, the younger substrate of the Weichselian glaciation is richer in nutrients. Field investigations were partly carried out ahead site destruction by opencast lignite mining (Berg-Hobohm 2004, Medović 2004) [31,32] or expansion of the airport Berlin-Brandenburg (Stika 2022) [33]. Göritz is situated on substrate of the Saalian glaciation (LBGR) [34]. It represents a late Roman period settlement continuously inhabited for more than 200 years since the mid of the 3^rd^ century. Some 150 houses were recovered and rich finds of rye grains and cattle bones were made (Berg-Hobohm 2004) [31]. The weed assemblage suggests a regular and intensive cultivation, that probably included crop rotation with fallow land, and thus some kind of manuring by grazing animals (Jahns et al. 2018) [30]. The rye from Groß Lübbenau is part of a rye dominated cereal mass find from a Slavic castle of the 9th/10th century, charred during a major fire that destroyed the castle (Medović 2004) [32]. In the high medieval Anger village Diepensee (13^th^ century to latest 1375 CE) rye was dominating as well. The occurrence of bread wheat (*Triticum aestivum*), the associated weeds and the poor record of millet (*Panicum miliaceum*) point to relatively good soils (Jahns et al. 2018, 2022, Stika 2022) [30,33,35]. The soil substrate around Diepensee is built of glacial sediments of the Weichselian (LBGR) [34]. Archaeozoological and genetic studies have proven the great importance of horse husbandry (Hanik et al. 2022) [36]. At the Slavic Dyrotz (late 10^th^/early 11^th^ to early 13^th^ century), located also in an area with Weichselian deposits, findings of rye are rare, while wheat was quite common (Jahns et al. 2018; Mackowiak 2011; Kennecke 2008; LBGR) [30,37,38]. For a better understanding of the isotope values, we compare them with that of Emmer (*Triticum dicoccum*) from the 6^th^ millennium BC of the Linearbandkeramik (LBK) period recorded from a pit at Lietzow (Kirleis et al. 2019, 2024) [39,40]. The geological ground consists of Weichselian deposits with peats in a moist depression nearby (LBGR).

Rather early rye co-dominated findings are those from the Dariali Fort in Georgia. Remains from the earliest occupation phase, radiocarbon-dated to mid-4^th^ to early 5^th^ century were analysed. Situated at 1350 m asl on a steep valley slope in the Caucasus, the Dariali Fort controls a pass from south-west Asia to eastern Europe in the north. Late medieval terraces point to local cereal cultivation, evidenced by archaeobotanical findings made up to 1950 m asl in the Caucasus. Abundance of animal bones in the Dariali Fort indicate animal husbandry and the availability of dung for agricultural practices (Shumilovskikh and Poole 2020) [41].

## Laboratory Methods

The charred archaeobotanical grains were cleaned with 6M HCl overnight to remove mineral adherents, washed and stored overnight in demineralized water. For Oldorf with its calcareous layers of marine clay, an additional subsample was treated with demineralized water only and measured for *δ*^13^C and *δ* ^15^N. For the *δ*^13^C the ANOVA test for equal means with the software Past v. 4.11 (Hammer and Ryan 2001) [42] indicate statistical significant differences (p < 0,01) and only the 6M HCl data are considered further, for the *δ*^15^N (p = 0,63) values of both treatments are combined. Additional subsamples excavated from the sandy soils of Neermoor were cleaned with 6M HCl only shortly, 10% HCl or demineralized water respectively. ANOVA tests for equal means pointed to no statistical significant differences (*δ*^13^C, p = 0,33; *δ* ^15^N, p= 0,40; *δ*^34^S, p = 0,26) and the results of the subsamples are combined here. The modern grains (Halle, Thyrow) were charred in open small Porcelain crucibles at 250°C for 5 h. Temperature of the muffle furnace was permanently controlled for its stability. All values in the text, figures and tables refer to charred grains without any disputable adjustments for charring (Kirleis et al. 2023) [43]. All grains were individually weighted, mortared and the powder weighted into tin capsules. For some measurements 2 to 10 grains were necessary to reach the needed sample weight.

The stable isotope laboratory of the GeoZentrum, Erlangen, employed a Flash EA 2000 elemental analyser connected online to a ThermoFinnigan Delta V Plus mass spectrometer to measure the *δ*^13^C and *δ*^15^N. Additional samples were permitted at the Centre for the Isotope Sciences, Glasgow, for analysis of *δ*^34^S, *δ*^13^C and *δ*^15^N using a Thermo Fisher Scientific Delta V Plus IRMS with an IsoLink Flash HT element analyser. Results are calibrated against the standards USGS 40, 41, MSAG2, M2, SAAG2 and reported as ‰ relative to VPDB for carbon, atmospheric air for nitrogen and VCDT for sulphur. In the text <u>mean values</u> are underlined, outliers (outside inner quartile range plus/minus 1.5 SD) are given in brackets.

The AMS ^14^C-results of three individual grains dated by the Radiocarbon Laboratory Poznań, Poland, are calibrated online using OxCal version 4.4 with the IntCal20 calibration curve (Reimer et al. 2020; Bronk 2009) [44,45].

## Results and discussion

The unmanured rye from the sandy soils of Thyrow exhibit *δ*^15^N values between -1.2 to 1.2 ‰ (<u>-0.5</u> ‰), what is close to the 0.0 ‰ *δ*^15^N of the atmosphere (Fig. 2). Therefore, N of rye might originate initially, beside others, from nitrogen fixation by Cyanobacteria or symbiotic nodule bacteria of legumes (Wada et al. 2012; Skrzypek et al. 2015) [46,47]. The manured rye (15 t dung /ha*a) shows much higher *δ*^15^N values of 4.5 to 8.0 ‰ (<u>6.3</u> ‰). This common pre-industrial level of manure application causes a clear increase in *δ*^15^N by some 7 ‰ in rye on sandy soils.

**Fig. 2.**
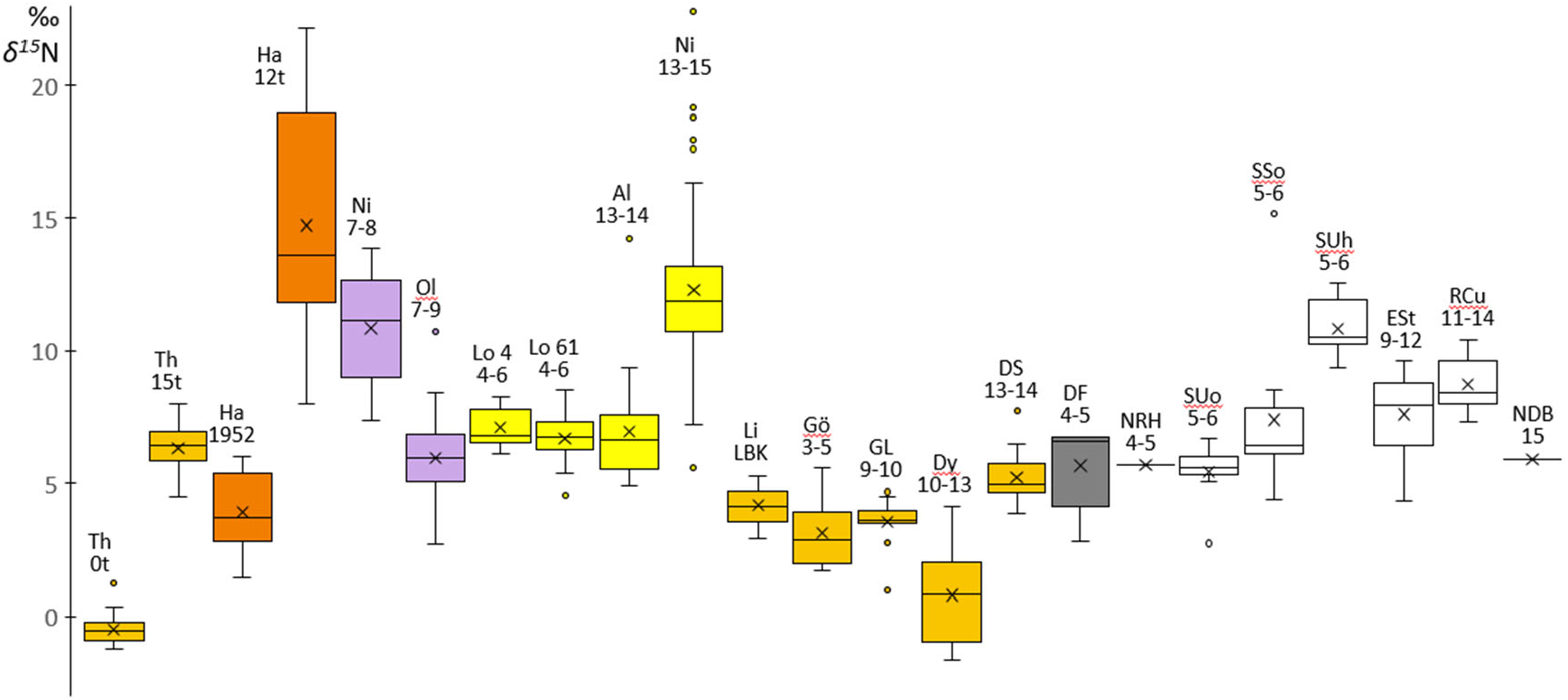
*δ*^15^N box plots of experimental sites (Th, Ha), Wurten (Ni, Ol), Geest (Lo, Al), eastern Germany (Li to Dy), Dariali Fort (DF) in Georgia, and from literature from the Netherlands (Netherlands, Raalte-Heeten, NRH, Deventer Burseplein, NDB), Sweden (Uppåkra oven, SUo, Stanstorp oven, SSo, Uppåkra house 11, SUh), England (Stafford, ESt), Romania (Cuneşti, RCu) (Brinkkemper et al. 2017, Larsson et al. 2019, Hamerow et al. 2020, Golea et al. 2023) [12–15]. For Th and Ha the numbers refer to the amount (t/ha*a) and last year of manuring respectively the expected age of the archaeological material (centuries CE); LBK, Linearbandkeramik.

On the loess soils of Halle, the rye manured from 1878 until 1952 (12 t dung/ha*a) shows values from 1.5 to 6.0 ‰ (<u>3.9</u> ‰), close below the *δ*^15^N level of the manured Thyrow experiment. The highest, but also most variable, *δ*^15^N exhibit the rye manured continuously since 1878 with values of 8.0 to 22.1 ‰ (<u>14.7</u> ‰) showing the high influence of 65 years of further manuring. Compared to the lower values from the stronger manured plot of Thyrow, a high positive influence of loess as initial soil substrate on the *δ*^15^N seems verified.

By combining *δ*^15^N results from agricultural experiments in northern Europe on different soils, using different cereals and different kinds of manure (Fraser et al. 2011) [48], Bogaard et al. (2013) [49] developed a widely used scale for manure intensity. In this standard, *δ*^15^N values below 3 ‰ stand for no, low or former manuring, values up to about 6 ‰ for medium (10-15 t/ha*a) and above 6 ‰ for high (35 t/ha*a) manure application. Following this scale, rye from Halle (12t/ha*a) is medium manured but delivers a *δ*^15^N (8 to 22.1 ‰) classified as highly manured. On the other hand, the slightly stronger manured rye from the sandy soils of Thyrow (15 t/ha*a), shows a much lower *δ*^15^N (4.5 to 8 ‰). That less manured rye from the loess soils shows a much higher *δ*^15^N may indicate that manuring can generate significant different isotope results depending on soil quality.

The earliest isotope data of rye for northern Germany is from the Geest site Loxstedt (4^th^-6^th^ century CE). The *δ*^15^N of the two samples representing different contexts range from (4.6) 5.4 to 8.5 ‰ with mean values of <u>6.7</u> ‰ respectively <u>7.1</u> ‰. The *δ*^15^N of the much younger rye of Altenwalde (13^th^-14^th^ century) ranging from 4.9 to 9.4 ‰ (to 14.2 ‰, <u>6.9</u> ‰) is close to the rye of Loxstedt – altogether much higher than for the unmanured rye from Thyrow, but close to the rye from Halle manured by 15 t dung /ha*a. Therefore, it seems, that already since the migration period or shortly after, intensive manuring of rye took place, long before the plaggen soil system started in high medieval times. The slightly wider *δ*^15^N range in Altenwalde may illustrate, that grains originated from more than one yield, thus from different fields or years.

The rye of the Wurt Oldorf (9^th^ century) falls into and below the *δ*^15^N range of the Geest whereas the values at Niens (7^th^-8^th^ century) exhibit with a range of 7.4 to 13.9 ‰ (<u>10.8</u> ‰) a much higher ^15^N content. As the dwelling mounds were partly built with stable dung, the high *δ*^15^N values of Niens may point to cultivation of rye on plots directly on the mounds. In addition to dung, the *δ*^15^N could partly originate from material collected from drift lines (Gröcke et al. 2021) [50]. The flooding by storm surges in winter makes it likely, that rye was cultivated in summer after rain has diluted the salt influence which has been still relatively low in the early settlement phase (Behre 1991) [24]. The *δ*^15^N of Neermoor, at the transition from the marshland to the Geest, displays the highest and most divers ^15^N enrichment of the archaeological remains ((5.6) 7.2 to 16.3 ‰ (22.8 ‰), <u>12,3</u> ‰) the highest. All values give evidence of soils rich in ^15^N, comparable to the contents at Niens or even higher. Including the upper outliers, the range covers nearly that of the manured loess-soils of Halle. Stable dung will probably have been a prominent ^15^N source, however, as discussed below, the *δ*^34^S points to an additional source.

The archaeobotanical rye from east of the river Elbe exhibits *δ*^15^N values clearly higher than the unmanured modern rye from Thyrow. The ^15^N enrichment of the medieval manuring at Diepensee (3.9 to 7.7 ‰) was already equivalent to about 15 t/ha*a stable dung. Parts of the rye from Göritz (1.7 to 5.6 ‰) points to quite intensive manuring already 1500 years ago as well as the rye from Groß Lübbenau (1.0 to 4.7 ‰) of the following Slavic period. Nevertheless, their *δ*^15^N level was already achieved by emmer in the Neolithic site Lietzow (2.9 to 5.3 ‰). Compared to the Thyrow experiments, at least some of the rye of Dyrotz (−1.6 to 4.1 ‰) received dung, while a certain part appears not manured. Low to negative *δ*^15^N values as at Dyrotz are reported for herbs and grasses growing on forest floors, due to ^15^N depleted tree litter (Sykut et al. 2021) [51]. Possibly the low *δ*^15^N-rye was cultivated on fields not manured since forest clearing and therefore reflecting modern clearings or fields located too far from the settlement for intensive managing.

At Oldorf, Altenwalde and Diepensee, single grains show high values beyond all others. Such grains are also reported by Larsson et al. (2019) and appear in the unmanured rye of Thyrow. These statistical outliers may originate from plants growing on extreme ^15^N-rich places, like close to dung heaps or on spots where domestic animals during stubble pasture or wild animals dropped on the fields.

Beside nutrients, the water availability controls the amount of grain yield. Under humid conditions with high yields, wide open stomata increase the CO_2_ inflow into the plants, leading to a higher rate in carbon fixation and starch synthesis. The high CO_2_ exchange with the atmosphere leads to a lower ^13^C content respectively a higher Δ^13^C in the grains. During water deficit a reduced fractionation effect leads to a lower Δ^13^C value. Thereby, the ^13^C content of cereal grains is a proxy for the hydrological conditions during the summer months, when the grains were built, and closely related to yield amounts. The conversion to Δ^13^C values calculates out the overtime changing atmospheric ^13^C content (Araus et al. 2003; Ferrio et al. 2005) [52,53]. Araus et al (2003) [53] developed a ^13^C based calculation of yields from archaeological wheat and barley grains for the Mediterranean region. Absolute amounts need, beside others, assumptions about former cultivation practices and yields of ancient varieties. We therefore calculate relative yield amounts for comparing rye originating from neighbouring former settlements with comparable agricultural development status. As formulas for absolute yield amounts, as developed by Araus et al. (2003) [53], need several arguable assumptions, we present here a formula for relative yields (ry). It is based on water deficit experiments with rye on loamy sands in north-eastern Germany and Poland (Kottmann et al. 2014, 2016; Schittenhelm et al. 2014) [54–56]. The results imply that a water shortage accounting for a Δ^13^C decrease of 1 ‰ results in a yield reduction of about 12.2 ‰. The yield reductions due to water stress reported by Kottmann et al. (2014) [54] is up to 60% and by Krafft (1918) [57], including different soils, of ca. 80%. The absolute grain yields of winter-rye on sandy soils in north-eastern Germany was around 1 t/ha in the first half of the 19^th^ century (Miedaner 2014) [58]. Körnicke (1885) [59] reports averages of 1 t/ha for whole Germany, 0,5 t/ha for the worse and 2 t/ha for the best fields. The yield of the manured rye of Halle was around 2.7 t/ha at the turn from the 19^th^ to 20^th^ century (Herbst et al. 2017) [20].

Based on mean Δ^13^C, rye yields were highest on the fields managed by the people of Niens (100 % ry), closely followed by Groß Lübbenau (93% ry), Oldorf (92 % ry), Altenwalde (88% ry) and Dyrotz (84 % ry) (Fig. 3). All other sites have less to much lower Δ^13^C values and therefore lower relative yields. Δ^13^C from the Caucasus points to a yield of 1/3 compared to Niens. The chain of effects from intensity of manuring to increased grain *δ*^15^N and higher grain yield is not understood yet. It can only be speculated, that the more intensively manured rye from Neermoor produced a yield close to that at Altenwalde (both on the Geest) or even higher. For the same reason the fields of Altenwalde might have been more productive than that of Groß Lübbenau. For a more detailed estimation of archaeological cereal yields in relation to water and nitrogen supply, dedicated experimental plots are required. Another variable but hard to determine, is a possible difference in yields between summer- and winter-rye, that was about 25% in the 19^th^ century (Körnicke 1885) [59].

**Fig. 3.**
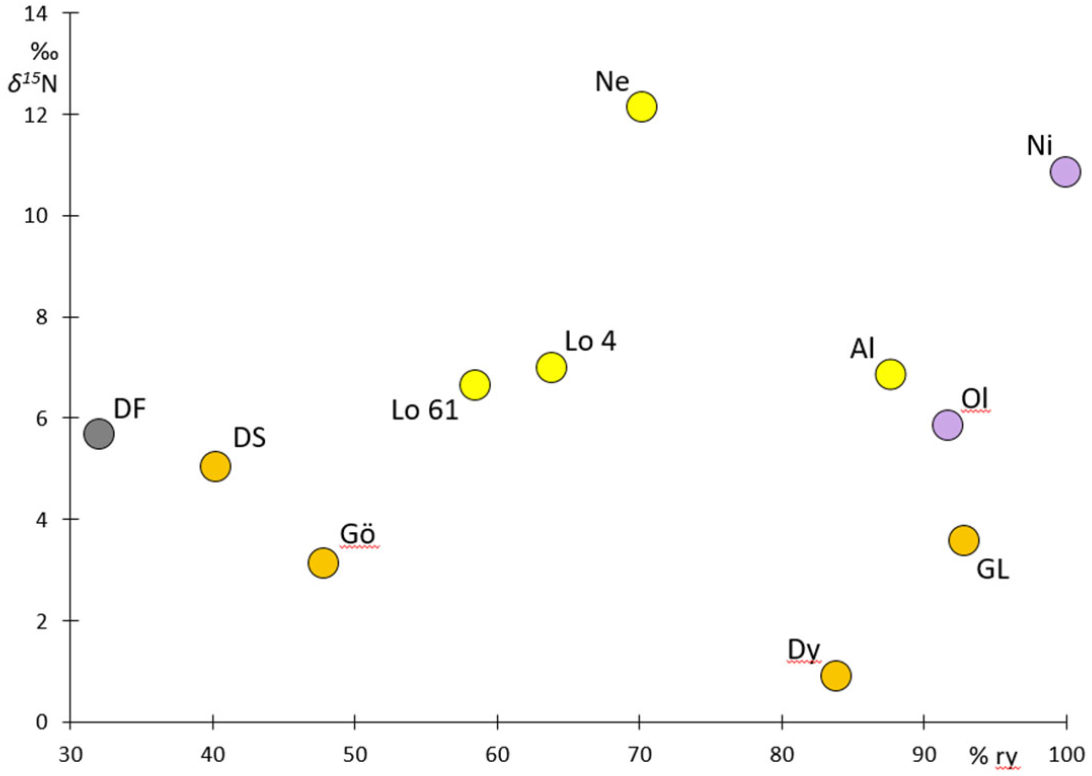
Mean relative yields and *δ*^15^N (acronyms as in Figs. 1 and 2).

The soil *δ*^34^S is controlled by geology and hydrology. Under wet, oxygen deficit conditions microbial processes strongly affect the sulphur isotope composition (Stack et al. 2011) [60]. Sulphur uptake happens nearly without fractionation and plants reflect quite closely the soil *δ*^34^S. It can range from about -5 to 22 ‰ in terrestrial plants, while *δ*^34^S of the ocean water is constant at 21‰ (Fry 2008, Tcherkez and Tea 2013) [61,62]. Reports on ^34^S from rye are scarce in literature respectively the data published here seems to be the first ones on archaeobotanical material. Due to quantities needed, measurements per site are very limited and therefore merged here for the environments Wurten (Oldorf, Niens), Geest (Altenwalde, Loxstedt), and eastern Germany (Göritz, Diepensee, Dyrotz). Neermoor stands for itself through many measures. To understand the possible palaeoecological significance of *δ*^34^S values, we also include data on emmer (*Triticum dicoccum*) from the Neolithic Lietzow site. With ranges of 8.9 to 14.7 ‰ and 6.6 to 11.2 ‰ the *δ*^34^S of rye from the Geest sites Altenwalde and Loxstedt and eastern Germany is quite similar, probably related to having in principal the same geological substrate as well as low water saturation of the porous soils (Fig. 4). The somewhat higher Geest values might originate from marine influence via sea spray (*δ*^34^S ca. 21 ‰) and marine precipitation (*δ*^34^S ca. 13 ‰) (Fry 2008) [61]. Although the Wurten are closer to the sea, the rye *δ*^34^S is only 1 to 10.4 ‰. The lower values might be associated with the different substrate and thus be controlled by the dung of the Wurten. With a *δ*^34^S of 2.3 to 8.4 ‰ one part of the rye from Neermoor falls within the same low range of the Wurten. A δ^15^N of 10.9 to 19.3 ‰ indicates, that the isotopes of these grains derive most probably from dung.

**Fig. 4.**
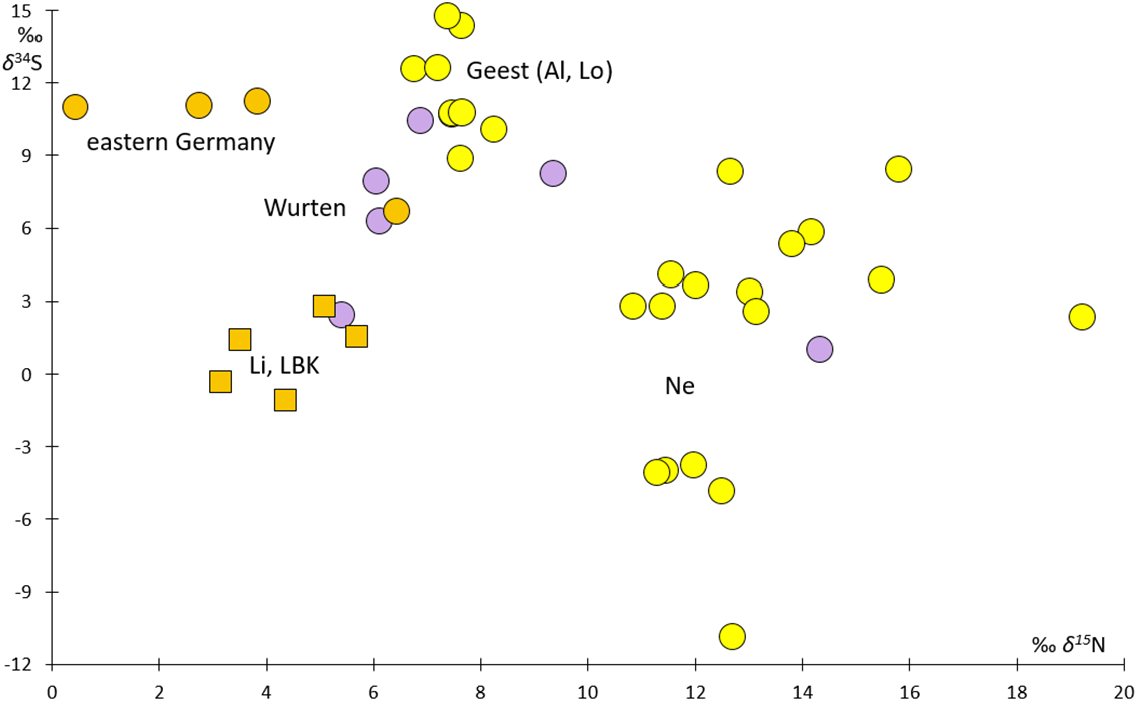
*δ*^34^S and d^15^N of individual grains grouped by region and for distinctive sites (acronyms as in Figs. 1 and 2).

The remaining grains of Neermoor exhibit also relatively high but small *δ*^15^N (11.3 to 12.7 ‰), but exhibit negative *δ*^34^S of -3.8 to -11 ‰. The peats near Neermoor consist of fen and raised bog peats on top and became flooded by salt water with rising sea level, what enriched them with salts including sulphates. A microbial fractionation during sulphate reduction under the anaerobic conditions of the peats can lead to a *δ*^34^S of around -10‰ or even -20‰ (Gerdes et al. 2003) [63] [61]. The cutting of such peats for salt extraction became widespread since the 12^th^ century at the latest (Behre 2008) [64]. The use of fen peat layers not suitable for the extraction of salt could be a reason of the low *δ*^34^S of the rye. Nevertheless, it can be expected that also the dung accumulating over winter in stables or dung heaps was at least in its deeper parts saturated by animal urine and thereby anaerobe, leading to *δ*^34^S reduction, as some grains from Niens and Oldorf suggest.

The Neolithic emmer from Lietzow also shows a striking low *δ*^34^S (−1.1 to 2.8 ‰) especially in comparison with rye from that area. The geological substrates have the same sources, but at Lietzow there is also a swampy area. Possibly, peat from the swamp was used for soil improvement of the fields, leading to the relatively low *δ*^34^S in the cereals. More focus on *δ*^34^S is needed to proof the above assumptions setting charred grains as archaeological and palaeoecological archive better in value.

## Discussion

In SW-Asia today the many varieties of weedy rye (subsumed in *Secale cereale* ssp. *ancestrale* Zhuk., Feldman and Levy 2023) [65] vary in their rachis traits from fully scattering over half scattering to non-scattering. As long as mechanical weed control is weak, scattering has advantages. Grains falling to the ground can sprout and establish new plants before the sowing of the cereals. This applies even more for winter forms having a start directly after the harvest making them strong competitors against later sown cereals, especially against the summer cereals of the following year. With more effective weed control by ploughing with the mouldboard plough this ahead strategy became a drawback. Non-scattering forms of weedy rye harvested, threshed and sown together with the cereals after ploughing became a positive selection advantage now. Thus, with the dissemination of the mouldboard plough in the last centuries BCE (Both et al. 2010) [66] weedy rye may have turned into a fully domestic weed with stiffy ears, fully depending on human threshing and sowing. Where drought and cold events periodically damage the cultivated cereals, farmers tolerate upcoming weedy rye that can make up to over 40% of the grain yield in adverse years (Hillman 1978) [1]. Depending on the frequency of dry years, the step to an intentional cultivation of rye might take quite a short time, particularly on sandy soils susceptible to dryness. In Brandenburg, an experimental field cultivated with a half and half mixture of wheat and rye turned by sowing from the harvest admixture into a nearly pure rye field after three years (Behre 1992) [3]. Therefore, it seems, beside other factors (Behre 1992) [3], that especially the intervention of the mouldboard plough on sandy soils susceptible to dryness, to the east also the cold-dry continental climate, may have turned weedy rye into a valuable staple cereal (*Secale cereale* ssp. *cereale*) in its own right.

In north-western Germany the level of cultivation was reached latest at the turn from the 1^st^ to the 2^nd^ century CE (Behre 1992, 2001, 2010) [3,67,68]. Due to the accompanying weeds, it seems that firstly summer-rye was cultivated, with the Geest around Altenwalde and Loxstedt as a centre of cultivation already in the Roman period (Behre 2001) [67]. It was also in the same period, that rye became a co-dominating cereal in Brandenburg (Jahns et al. 2018) [30]. The turn to winter-rye took place at the beginning of the first millennium.

Rye finds occur occasionally on the Wurten along the North Sea coast since the Iron Age, become more frequent after the Migration period and are interpreted to be an import from the Geest (Schmid 1994, Schepers and Behre 2023) [23,69]. The differences between the isotope signatures of the Wurten and the Geest contradict the assumption of an import, and summer-rye was more probably cultivated on the Wurten, as discussed above. Nevertheless, further analyses may provide indications also for imported cereals. The isotopic signals from Niens but even more the early manuring at Loxstedt as well as Göritz seem to contradict the idea of cultivating rye because of its frugality.

Some kind of manuring is clearly needed to maintain higher yields not only on sandy soils, where nitrogen is the most limiting nutrient for rye production (Ellmer et al. 2005) [70]. Even on the fertile loess soils of the Eternal Rye experiment of Halle started in 1878, the decadal mean yield from the unmanured rye plot halved within the first four decades from 2,3 t/ha to 1,1 t/ha (Herbst et al. 2017) [20]. Compared to crop systems with fallow years for some recovery, the use of manure may be much more effective, but depends on manure availability and knowledge. As evidenced by house floor plans from the coast area, stables exist there as in Scandinavia since the Bronze Age (Behre 2002, Zimmermann 1999, Zimmermann 2014) [71–73], and dung could also have been collected for instance from pastures or kraals (Schlütz et a. 2023; Zimmermann 1999) [72,74].

In the sandy lowlands facing to the North Sea, the application of dung, settlement waste and heath sods to increase for instance the fertility of Celtic fields dates back already to the first millennium BC (Blume and Kalk 1986, Blume and Leinweber 2004, Arnoldussen 2018) [75–77]. South of the Baltic Sea the increase of soil quality by human management may have started around the turn from Bronze to Iron Age (Acksel et al. 2017) [78], and there is evidence of anthropogenic improvement of soil fertility by settlement waste and possibly dung for Brandenburg since the 7th century BC (Neef et al. 2002) [79].

Obviously, dung availability and the knowledge of its wise use was wide spread and practiced in northern Europe before rye cultivation. Under certain circumstances it was obviously a big advantage to manure even the undemanding rye. Due to the poor soils, the Geest was occupied in the Neolithic only by single farmsteads (Zimmermann 1995) [80]. Increased yields through manuring probably enabled the transition to villages with larger populations, taking place on the Geest during Roman times (Zimmermann 1995) [80]. At some places with wheat cultivation, like possibly in Diepensee, rye may have grown on manured fields in crop rotation with the demanding wheat. In case of Dyrotz, rye was cultivated on poor soils, possibly not suitable for wheat. The comparably higher rye *δ*^15^N on the Geest sands might point to an ample access to dung by the extensive cattle breeding on the fat pastures in marshland in front (Schmid 1994) [23].

On a spatio-temporal perspective, the manuring level of rye fields does show no trend during time. Even within the oldest periods covered, the samples of Göritz (1^st^-4^th^ century), Georgia (4^th^century), Loxstedt (4^th^-6^th^ century) and Niens (7^th^-8^th^ century) represent all levels from low to intensive manuring. This applies even more for the medieval samples, having the lowest (Dyrotz, 10^th^-13^th^ century) to the highest (Neermoor, 13^th^-15^th^ century) ^15^N content of all samples (Fig. 2). The mean *δ*^15^N values from rye outside Germany stretch over a range from 5.7 to 11.4 ‰. Most of the sample averages and ranges are around the values measured at Thyrow (application of 15 t dung/ha*a). This includes the oldest material from the 4^th^ to 5^th^ century from Georgia as well as the youngest sample Deventer Burseplei from the 15^th^ century of the Netherlands (Brinkkemper et al. 2017) [13]. Obviously, rye from a fertile loess area in Romania (11-14^th^ century) (Golea et al. 2023) [15]as well as from the sample Upper House (5-6^th^ century) on the poor soils of Sweden (Larsson 2019) [12] have the highest ^15^N contents. Their *δ*^15^N values fall completely or partially into the lower range of the loess soils of Halle (12 t/ha*a). The rye from Uppåkra/house 11 (9.4 to 12.6 ‰) was nearly as intensive manured as the rye of Niens and Neermoor.

In contrast, the close by find Uppåkra/oven exhibit with 2.7 to 6.7 ‰ a significant lower ^15^N level. Obviously, the people of the regional centre Uppåkra consumed rye produced under very different manuring regimes. While Kanstrup et al. (2014) [81] found a long-term trend in cereal manuring over the millennia BCE, there is obvious no time trend in archaeobotanical rye on any level from within or between regions of German nor throughout Europe. The trend to higher values at the German coast, as compared to the inland, is most probably caused by the plenty of dung from the high number livestock population on the fat meadows of the German clay district.

## Conclusion

Isotope analyses ((*δ*^13^C, *δ*^15^N, *δ*^34^S)) indicate that cultivation of rye in northern Europe was promoted by substantial manuring at the latest since the Migration Period. The transformation of rye from a weed to a domesticated crop was apparently fosteredby a number of technological developments (Behre 1992) [3]. In particular, the intensification of weed control through the introduction of the mouldboard plough around the birth of Christ may have resulted in rye traits that were pre-adapted to domestication by the formation of non-shattering ears. Weedy rye may have served initially as a backup crop in years of harvest failure due to cold or drought, before recognition as a cereal in its own right. Its productivity was enhanced through the application of animal dung and other forms of organic matter on poor soils. It appears that the rise in rye production as a result of manuring may have contributed to the advent of social change and later reinforced social hierarchy and inequality towards the medieval period. The application of manure is thus one factor that contributed to the social transformation from the traditional individual farmsteads towards the emergence of multi-headed village communities and towns.

No discernible trend in manuring practices could be identified across the periods under consideration. It seems reasonable to posit that the utilisation of manure was an intentional decision depending on local requirements and opportunities. The current evidence does not support the hypothesis that rye was imported from the Geest to the marshlands. Instead, it is more plausible that rye was produced locally on the elevated dwelling mounds in the marshlands

The relatively high ^15^N level of the dwelling mounds and the Geest is associated with the high cattle population in the surrounding fertile marshland and the accumulation of manure from the winter-housing of the animals. In addition, peat was employed as fertiliser at Neermoor. The discovery of large quantities of rye in medieval castles, including Neermoor, Groß Lübbenau, the Slavic Starigard, Parchim and Tornow (Kroll and Willerding 2004, Alsleben 2012, Jäger 1966) [82–84], several churches on the Geest (Behre 1973, Kučan 1979) [85,86] and cities such as Bremen (Behre 1991) [87] indicates that the elites in northern Germany and in the Slavic area east of the Elbe maintained their authority by stockpiling rye. The transition from summer-rye to the permanent cultivation of winter-rye during the medieval period resulted in rye becoming the primary cereal crop until its replacement by wheat in the mid-twentieth century enabled by chemical fertiliser.

Further investigation may elucidate whether evidence of manuring can be proven for the process of domestication and earliest cultivation. Experimental rye cultivation may further elucidate whether rye is superior to other cereals with regard to manuring. Nevertheless, rye may have been the preferred cereal on poor soils, where manuring was not feasible due to a lack of dung or distances that were too great to allow for manuring to be carried out effectively (Dyrotz). In order to gain a clearer estimation of archaeological cereal yields in relation to water, nitrogen and soil quality, it would be beneficial to set up specific experimental plots. Furthermore, this could facilitate the generation of reliable estimates of absolute crop yields.

It is evident that *δ*^35^S is capable of differentiating between subgroups that cannot be distinguished by ^13^C and ^15^N (Neermoor). Furthermore, *δ*^35^S has the potential to identify processes related to hydrological processes in the widest sense, taking place in peats, dung and soils. This enables the disclosure of environmental conditions and cultivation techniques. The application of triple isotope analysis to a range of crops and weed species could enable the generation of a comprehensive understanding of agricultural management practices at multiple scales, including landscape, site, species, cultivation system and even single field levels.

## Acknowledgments

Research, writing and open access to this paper was made possible within the scope of the Cluster of Excellence ROOTS “Social, Environmental, and Cultural Connectivity in Past Societies” under Germany’s Excellence Strategy (EXC 2150 - 390870439) and the project “Kontinuitäten in Zeiten des Umbruchs – interdisziplinäre Untersuchungen zu Klima, Umwelt und kultureller Entwicklung im 1. Jahrtausend n. Chr, in Nordwestdeutschland” (74ZN1363, Continuities in times of upheaval - interdisciplinary studies on climate, environment and cultural development in the 1st millennium AD in Northwest Germany) at Kiel University and at the Niedersächsisches Institut für historische Küstenforschung (NIhK, Lower Saxony Institute for Coastal Research) in Wilhelmshaven, funded by the Deutsche Forschungsgemeinschaft (DFG, German Research Foundation) and by the Ministry for Science and Culture of Lower Saxony, Germany. Part of the field work was financed by the DFG (111676444) and the ERC Persia. The funding agencies had no role in the design, execution of the experiment nor in the data analysis and manuscript writing. We would like to thank Helmut Eißner, Halle/Saale, for his generous support with material.

